# Histidine limitation causes alteration in the TOR network and plant development

**DOI:** 10.1101/2024.06.17.599310

**Authors:** Amandine Guérin, Caroline Levasseur, Aline Herger, Dominik Renggli, Alexandros Georgios Sotiropoulos, Gabor Kadler, Xiaoyu Hou, Myriam Schaufelberger, Christian Meyer, Thomas Wicker, Laurent Bigler, Christoph Ringli

## Abstract

Plant growth depends on growth regulators, nutrient availability, and amino acids levels. The TOR (Target of Rapamycin) network senses these parameters and influences cell wall formation and expansion accordingly. Cell wall integrity and structures are surveyed and modified by a complex array of cell wall integrity sensors, including LRR-extensins (LRXs), that function as hormone receptors and help to compact cell walls. Expressing the Arabidopsis root-hair specific LRX1 without the extensin domain, which anchors the protein to the cell wall, has a negative effect on root hair development. The mechanism of this negative effect was investigated by a suppressor screen, which led to the identification of a *sune* (*suppressor of dominant-negative LRX1*) mutant collection. The *sune82* mutant was identified as an allele of *HISN2* which encodes an enzyme essential for histidine biosynthesis. The *sune82* mutation leads to reduced accumulation of histidine, and this influences the TOR network. The *sune82* mutant reflects the impact of the TOR network on cell wall formation processes involving LRX proteins. It also represents an excellent tool to study the effects of reduced histidine levels on plant development, as it is a rare example of a viable partial loss-of-function allele in an essential biosynthetic pathway.

**Highlight:** Partial loss of function of *HISN2* in *sune82* results in a significant reduction in histidine content, which subsequently alters the TOR network.

## INTRODUCTION

Growth and development of any given organism depend on the availability of nutrients and essential organic molecules such as vitamins or amino acids used for protein biosynthesis. Unlike many animals including humans, plants are able to synthesize all amino acids (Trovato *et al*., 2021). The homeostasis of the different amino acids is essential for and depends on the growth and metabolic activity of the organism, since amino acids are not only used for protein translation, but they also serve as signals, precursors for secondary metabolites and, in the case of BCAAs (branched chain amino acids) and Lys, as a carbon source for respiration (Heinemann and Hildebrandt, 2021). The biosynthesis of amino acids is tightly controlled, and their availability in turn influences cellular activities. The TOR (target of rapamycin) network is a key regulator of cell growth in eukaryotes (Loewith, 2011; McCready *et al*., 2020). In response to growth factors and amino acids, TOR regulates metabolic and translational activities, as well as the dynamics of cellular processes such as cytoskeletal organization (Dobrenel *et al*., 2016*a*; González and Hall, 2017; Cao *et al*., 2019; Lutt and Brunkard, 2022). Central to this pathway is the TOR kinase, a phosphatidylinositol 3-kinase-related protein kinase that forms a TOR complex with interacting proteins such as RAPTOR and LST8 (Harris and Lawrence, 2003; Moreau *et al*., 2012). Several downstream targets of the TOR kinase have been identified, one of the best characterized being the S6 kinase, which phosphorylates the ribosomal protein RPS6 to influence ribosomal activities (Xiong and Sheen, 2015; Dobrenel *et al*., 2016*b*). The TOR kinase can be specifically inhibited by rapamycin, which was instrumental in the identification of the TOR kinase (Chan *et al*., 2000), but also by a new generation of inhibitors, including AZD-8055, which are valuable in modifying the TOR network (Montané and Menand, 2013). A number of proteins involved in the TOR network have been identified in various organisms, including plants, based on the observation of altered sensitivity to TOR kinase inhibitors (Leiber *et al*., 2010; Barrada *et al*., 2019; Cao *et al*., 2019; Schaufelberger *et al*., 2019; Chen *et al*., 2023).

An important regulator that directly interacts with the TOR kinase is FERONIA (FER), a CrRLKL-type transmembrane receptor protein that plays a central role in the control of a multitude of processes, including cell wall integrity sensing, cell growth, cell-cell recognition and cell fusion during fertilization, and immune responses (Escobar-Restrepo *et al*., 2007; Stegmann *et al*., 2017; Feng *et al*., 2018; Ortiz-Morea *et al*., 2021; Song *et al*., 2022). FER functions in conjunction with LRR-extensins (LRXs), extracellular proteins that serve as binding sites for RALF peptides that help restructure the cell wall matrix by inducing pectin compaction (Dünser *et al*., 2019; Herger *et al*., 2020; Moussu *et al*., 2020, 2023; Zhang *et al*., 2020; Schoenaers *et al*., 2024). LRX1 of Arabidopsis is expressed in root hairs and the defect in root hair development of the *lrx1* mutant is suppressed by altering the TOR network (Baumberger, 2001; Leiber *et al*., 2010; Schaufelberger *et al*., 2019). The suppressor of *lrx1*, *rol17* (*repressor of lrx1 17*) is mutated in the gene encoding IPMS1, an enzyme involved in leucine biosynthesis, resulting in increased valine levels (Schaufelberger *et al*., 2019). *rol17*, and another *IPMS1* allele, *eva1,* were found to alter the TOR network. This was evidenced by a reduction in sensitivity to the TOR inhibitor AZD-8055, which results in changes in cell growth and actin dynamics (Cao *et al*., 2019; Schaufelberger *et al*., 2019).

LRX proteins consist of an N-terminal LRR domain fused to an extensin domain (Herger *et al*., 2019). The extensin domain exhibits the typical characteristics of hydroxyproline-rich glycoproteins, which are structural proteins that can form intra- and intermolecular networks stabilized by covalent interactions (Mishler-Elmore *et al*., 2021). The extensin domain of LRX1 was demonstrated to insolubilize the protein in the cell wall matrix (Ringli, 2010). This domain is essential for LRX1 function, as expression of an LRX1 variant lacking the extensin domain, termed *LRX1ΔE14* (Ringli, 2010), fails to complement the *lrx1* mutant, and induces a root hair formation defect in wild-type plants. This dominant-negative effect is phenotypically stronger than the *lrx1* mutant (Baumberger, 2001).

In an alternative genetic approach to identify new factors influencing LRX-related processes, a suppressor screen was performed using the *LRX1ΔE14*-expressing line. This allowed for the identification of *sune* (*suppressor of dominant-negative LRX1ΔE14*) mutants, which alleviate the root hair formation defect observed in *LRX1ΔE14*. *sune82* was identified as an allele of *HISN2*, encoding an enzyme essential for histidine (His) biosynthesis. *sune82* contains a missense mutation that reduces His levels to a degree that is tolerable for the plant. The *sune82* mutation also affects the sensitivity to the TOR inhibitor AZD-8055, indicating that the TOR network is impacted in this mutant. The *sune* mutant collection was then screened for other genes that are known to affect TOR activity, leading to the identification of a new *rol17* allele, *sune106*. The identification of the *sune* mutants demonstrates that interfering with the biosynthesis of different amino acids, including non-BCAAs such as His, can affect the TOR network and alter cell growth processes.

## MATERIALS AND METHODS

### Plant growth and propagation

The *LRX1::LRX1ΔE14* line (referred to as *LRX1ΔE14* line) is in the Columbia background and described (Ringli, 2010). Unless otherwise described, the plants were grown on ½ MS/Vitamins with 2% sucrose, 10 mg/L myo-inositol, 0.5 g/L MES pH 5.7, and 0.6% Gelrite (Duchefa) in vertical orientation at 22°C and 16 h light: 8 h dark cycle. For propagation and crossing, seedlings were put in soil and grown under the same temperature and light regime. For the AZD-8055 (Chemdea CD0348) treatment, lines were germinated and grown on either DMSO or 0.2/0.4/0.6 μM AZD-8055 diluted in DMSO. The AZD-8055 stock solution was diluted so that an equal volume of treatment was added to the MS medium for the different concentrations.

For the histidine supplementation, L-Histidine (Sigma Aldrich H8000) was dissolved in water to prepare a 100 mM stock. Seeds were directly germinated on MS medium with or without 100 μM His.

For the phosphorylation assay, seeds were germinated on ½ MS/Vitamins with 0.3% sucrose, 0.5 g/L MES pH 5.7, and 0.5% Phytagel. 6-day-old seedlings were transferred to ½ MS liquid sucrose-free medium for 24 h and then either mock- or sucrose-treated (0.5%) for 4 h.

### EMS mutagenesis

The EMS (ethyl methanesulfonate) mutagenesis was performed similarly to Kim *et al*. (2006). Seeds from a wild-type Columbia plant expressing *LRX1::LRX1ΔE14* were incubated in 100 mM phosphate buffer overnight. The next day, seeds were incubated in 100 mM phosphate buffer containing 0.2% EMS for 8 h on a shaker. The M1 seeds were rinsed 15 times with 300 mL of water and then grown directly on soil in 240 pots, each containing 10 plants, to propagate to the M2 generation. Each pot represents a batch of M2 seeds. On average, around 20 seeds per M2 plant (200 seeds per batch) were screened for seedlings with a suppressed *LRX1ΔE14* root-hair phenotype. Selected putative mutant M2 seedlings were propagated and confirmed in the M3 generation. Positive lines were crossed with the non-mutagenized parental *LRX1ΔE14* line and propagated to the F2 generation that was analyzed for segregation of the *sune* mutant phenotypes.

### Whole genome sequencing, CAPS marker design, and targeted sequencing

Ten F2 seedlings from the first backcross of *sune82* with the parental *LRX1ΔE14* line exhibiting a *sune82* phenotype were selected and pooled for DNA extraction. Whole genome sequencing of *sune82* DNA, along with the non-mutagenized *LRX1ΔE14* line, was performed at Novogene using Illumina short read technology. Raw sequence reads from the pooled *sune82* mutants were trimmed with the Trimmomatic (version 0.38) with the parameters LEADING:10, TRAILING:10 SLIDINGWINDOW:5:10, MINLEN:50. Trimmed sequence reads were mapped with the bwa software (version 0.7.17-r118) to the Arabidopsis Columbia reference genome (TAIR version 10) using default parameters. Resulting Bam files were sorted and duplicates removed with samtools (version 1.9). New read groups were assigned to the reads with the Picard software (version:2.27.5). Sequence variants were called with the GATK software (version 4.2). The vcftools software (version 0.1.16) was used to filter the vcf files using the parameters (--max-meanDP 7 –remove-indels). The analysis revealed three SNPs (simple nucleotide polymorphisms) on chromosome 1 linked with the *sune82* mutation. Using this information, CAPS (cleaved amplified polymorphic sequences) markers were established, and co-segregation of the SNPs with the *sune82* phenotype was analyzed.

For sequencing of the *LRX1ΔE14* construct in the identified *sune* mutants, the construct was amplified using LRX1_F1 and LRX1_TermR primers (Table S1), targeting the promoter and terminator of *LRX1*, respectively. Due to the repetitive nature and length of the extensin coding sequence, only the *LRX1ΔE14* construct was analyzed and amplification of the endogenous *LRX1* was not possible.

For selection of plants lacking *LRX1ΔE14*, PCR to detect *LRX1ΔE14* with the primers LRX1_F1 and LRX1_TermR (see above) was conducted. To confirm the selection, seeds produced by PCR-negative plants were then grown on kanamycin, resistance to which is conferred by the *LRX1ΔE14*-containing transgene (Ringli, 2010 Ref).

The *IPMS1* gene in *sune10* and *sune65* was amplified by PCR and sequenced using different primers flanking and distributed in the coding sequence.

### CRISPR/Cas9-induced mutagenesis

The *LRX1ΔE14* parental line was transformed using a guide RNA targeting the gene of interest (Suppl. Table1) designed using CHOPCHOP (https://chopchop.cbu.uib.no/) and inserted into the *pAGM55261* vector (Grützner *et al*., 2021). T1 plants were selected based on RFP fluorescence, grown in soil, and the target gene was PCR-amplified and sequenced. T1 plants that were heterozygous or homozygous for *CRISPR/Cas9*-induced mutations were propagated to the next generation. RFP-negative (to select against the continuous presence of the guide RNA-containing pAGM55261) T2 plants were grown and again analyzed for the *CRISPR/Cas9*-induced mutations.

### Quantification of His levels

Plants used for His quantification were grown under standard conditions for 10 days. Whole seedlings were ground in liquid N2 and plant material was weighed and diluted with the following extraction buffer at 1:10 w/v: 80% methanol, 19% water, and 1% formic acid. The mixtures were vortexed for 5 s and placed in a sonic bath for 10 min. The mixtures were then vortexed again for 5 s and centrifuged for 5 min at 5000 rpm. The supernatant was transferred to an LC-MS vial. For His quantification, liquid chromatography was performed on a Thermo Fisher UltiMate 3000 UHPLC (Waltham, MA, USA). The UHPLC was built with a binary RS pump, an XRS open autosampler, and a temperature-controllable RS column compartment. Sample separation was performed at 30 °C using an HSS T3 Premier column (2.1 × 100 mm, 1.7 μm particle size) protected by the corresponding VanGuard pre-column (2.1 × 5 mm, 1.7 μm particle size) from Waters. The mobile phase consisted of eluent A (H2O + 0.02% TFA) and eluent B (MeCN + 0.02% TFA). The following gradient was applied at a constant flow rate of 350 μL/min: (i) 0% B isocratic from 0.0 to 0.8 min; (ii) linear increase to 90% B until 2.5 min; (iii) holding 90% B until 3 min; (iv) change until 3.1 min to the starting conditions of 0% B (v) equilibration for 1.9 min resulting in a total run time of 5 min. Mass spectra were acquired using a Thermo Fisher Scientific Q Exactive hybrid quadrupole-Orbitrap mass spectrometer (Waltham) equipped with a heated ESI source at position B and a voltage of 3.0 kV. Sheath, auxiliary, and sweep gas (N2) flow rates were fixed at 40, 15, and 1 (arbitrary units), respectively. The capillary temperature amounted to 275 °C, and the auxiliary gas heater temperature was 350 °C. The S-lens RF level was set to 55.0. The instrument was calibrated at a mass accuracy ≤ 2 ppm with a PierceTM LTQ Velos ESI Positive Ion Calibration Solution (Thermo). Data were acquired in full scan mode between *m/z* 50 and 750, at 70’000 full width at half maximum (FWMH), with resolution at *m/z* 200, a maximum IT of 247 ms, and an AGC target of 3e6. Xcalibur 4.1 and TraceFinder 4.1 softwares (Thermo Fisher Scientific) were employed for data acquisition, peak-area integration (extracted ion chromatograms with 3 ppm MS-peak width), and quantitation.

### Propidium iodide staining

Propidium staining was conducted by 2 min incubation of 5-day-old Arabidopsis seedlings in propidium iodide (Sigma Aldrich P4864) diluted to 1 µg/mL in water. Seedlings were then immediately washed for 3 min and mounted in water. A TCS SP5 laser scanning confocal microscope (Leica) with a 20x immersion objective (Leica 20x/0.75 NA) was used for imaging. Propidium iodide was excited at 561 nm with 30% laser power and detected at 610 to 620 nm. After acquisition, images were stitched together to obtain a complete image of the root. Cell lengths were measured manually in Fiji, starting from the beginning of the meristematic region.

### Immunoblotting

*LRX1ΔE14* harbours a cMyc tag at the beginning of the LRR domain, which does not affect protein function and enables protein detection by immunoblotting (Baumberger, 2001; Ringli, 2010). For the exact position of the cMyc tag, the sequence of LRX1cMyc is deposited as NCBI accession number GU235992. Root material of 100 seedlings grown for seven days on half-strength MS plates in a vertical orientation was collected, frozen in liquid nitrogen, and macerated with glass beads. Proteins were extracted using 100 µL of 1% SDS, 5 mM CaCl2, and cOmplete Protease Inhibitor Cocktail (Roche). After boiling for 5 minutes, samples were cooled on ice, centrifuged, and 20 µL of supernatant was mixed with 5 µL of 5x Lämmli buffer containing 5 mM DTT. The samples were boiled at 95 °C for 5 minutes, were then cooled on ice for 3 minutes, and subsequently centrifuged. 20 µL of protein extracts were loaded on a 10% SDS-PAGE gel using standard procedures. Following protein separation, blotting on a PVDF membrane was done using wet electroblotting (Bio-Rad). After blocking the membrane overnight with 1xTBS, 0.1% Tween 20, 5% low-fat milk powder, antibody incubation was done at 1:3000 dilution for both the primary anti-c-Myc (9E10, Thermofisher Scientific) and the anti-mouse IgG-HRP (Sigma Aldrich A4416) antibodies. Immunodetection was performed using Pierce™ ECL Western Blotting Substrate (Thermo Scientific) and a FUSION FX imager (Vilber) was used for blot visualization.

For the phosphorylation assay, whole seedlings were ground in Eppendorf tubes using a pellet pestle and total proteins were extracted in SDS extraction buffer (40 mM Tris-HCl pH 7.5, 10% glycerol, 5 mM MgCl2, 4% SDS, 1x protease inhibitor cocktail (Sigma-Aldrich), 1 mM phenylmethanesulfonyl fluoride). Protein concentrations were measured using the Pierce™ BCA Protein Assay Kit (Thermofisher scientific). 30 µg of total protein extracts were mixed with 1/5 volume of 5x Lämmli buffer (250 mM Tris-HCL pH 6.8, 5% SDS, 50% glycerol, 0.02% bromophenol blue), 5% β-mercaptoethanol. Proteins were separated on a 12% SDS-PAGE gel and transferred to PVDF membranes by wet electroblotting (Bio-Rad). To improve signal detection, membranes were treated with the Western blot enhancer SuperSignal™ (Pierce). Membranes were then blocked in 5% non-fat dry milk solution in PBS (137 mM NaCl, 2.7 mM KCl, 10 mM Na2HPO4, 2 mM KH2PO4, pH7.4) and then probed overnight with either mouse anti-RPS6 (dilution 1:1000, Cell Signaling), or rabbit anti P-RPS6 (dilution 1:3000) antibodies (Dobrenel *et al*., 2016*b*) at 4 °C. Goat anti-rabbit IgG-HRP (Sigma Aldrich A0545) and goat anti-mouse IgG-HRP (Sigma Aldrich A4416) were used as secondary antibodies (dilution 1:5000). Immunodetection was performed using Pierce™ ECL Western Blotting Substrate (Thermo Scientific) or, for lower signals, with SuperSignal™ West Femto Maximum Sensitivity Substrate (Thermo Scientific). A FUSION FX imager (Vilber) was used for blot visualization. Transferred proteins on PVDF membranes were visualized by Coomassie staining to check for equal loading.

### Statistical analyses

At least 5 biological replicates were used for each experiment. Graphs and statistics were generated using RStudio. A Shapiro-Wilk test was used to test the normality assumption of the data. If the normality assumption was not rejected, a Student t-test was used to compare the means of two given data sets. In this study, P-values< 0.05 were considered statistically significant.

## RESULTS

### *sune82* suppresses the dominant-negative effect of *LRX1ΔE14* on root hair development

As expression in wild-type Col of a truncated LRX1 lacking its extensin domain (*LRX1ΔE14*) leads to a dominant-negative effect (Baumberger, 2001; Ringli, 2010) (Fig. 1A, 1B), we took advantage of this striking root-hair phenotype in a genetic attempt to identify new components able to modify the LRX1-modified process. A transgenic line expressing *LRX1ΔE14* (Ringli, 2010) was used for EMS mutagenesis. Progenies of 1,000 M1 plants were screened for suppression of the *LRX1ΔE14*-induced root hair growth defect (for details, see Material and Methods). Twenty-two suppressors called *suppressor of dominant-negative effect* (*sune*) mutants were identified and one line, *sune82*, was analyzed in detail. This line, referred to as *LRX1ΔE14 sune82*, exhibits a partial suppression of the *LRX1ΔE14* root hair phenotype (Fig. 1A) with more and longer root hairs developing (Fig. 1B), but also displays a severe dwarf phenotype with shorter primary roots (Fig. 1C, 1D) and stunted adult plants (Supplementary Fig. S1) compared to *LRX1ΔE14*.

**Fig. 1.**
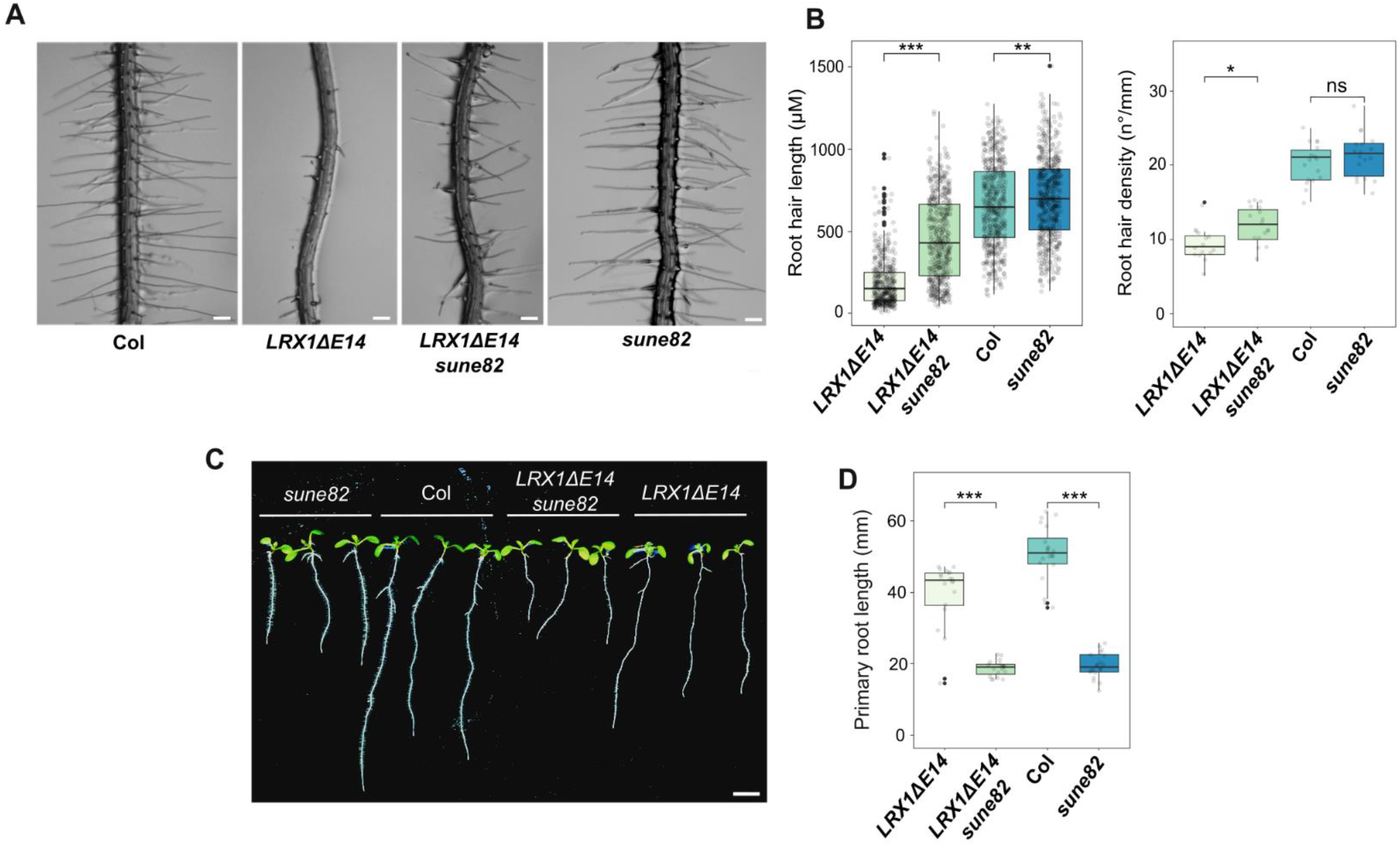
*sune82* suppresses the dominant-negative effect of *LRX1ΔE14* on root hair development. (A) 5-day-old roots of Col, *LRX1ΔE14*, *LRX1ΔE14 sune82,* and *sune82.* Seedlings were grown in a vertical orientation (scale bar 200 µm). (B) Quantification of root hair length (≥ 17 roots for each genotype, 30 root hairs were measured per root, n≥ 510) and root hair density (≥ 17 roots for each genotype, n≥ 17). (C) 10-day-old seedlings grown as in (A) (scale bar 5 mm). (D) Primary root length quantification of 10-day-old seedlings (≥ 20 roots for each genotype, n≥ 20). Asterisks on the graphs indicate significant differences between genotypes (*= P<0.05, **= P<0.01, ***= P<0.001) and ns indicate non-significant differences (P> 0.05) using an unpaired *t*-test. Black line in the boxplots represents the median.

Backcrossing of *LRX1ΔE14 sune82* with the parental *LRX1ΔE14* line produced F1 seedlings with an *LRX1ΔE14* phenotype, indicating that the *sune82* mutation is recessive. In the F2 generation only 15% of the seedlings showed a suppression of the *LRX1ΔE14* phenotype (1288 *LRX1ΔE14* and 188 *LRX1ΔE14 sune82* phenotypes). This segregation differs significantly from the expected Mendelian 3:1 segregation, indicating an impact of *sune82* on fertilization efficiency.

To determine the effect of the *sune82* mutation in the wild-type background, the *LRX1ΔE14 sune82* mutant was crossed with wild-type Col. In the F2 generation, plants with a short-root *sune82* mutant phenotype but lacking *LRX1ΔE14* were selected (see Material and Methods). *sune82* single mutants develop significantly longer root hairs than Col (Fig. 1A, 1B) and a primary root comparable to *LRX1ΔE14 sune82* (Fig. 1C, 1D). This, together with the stunted adult plant phenotype (Supplementary Fig. S1), reveals that the effect of the *sune82* mutation is widely involved in cell growth and does not solely influence the LRX1-related root hair developmental process.

### The *sune82* mutation affects the *HISN2* gene involved in His biosynthesis

To identify the *sune82* mutation, F2 plants of the backcross of *LRX1ΔE14 sune82* x *LRX1ΔE14* that showed a *sune82*-like phenotype were selected and pooled, and DNA was extracted for whole-genome sequencing, along with DNA of the non-mutagenized *LRX1ΔE14*. Homozygous mutations specific to the suppressor lines and absent from the parental *LRX1ΔE14* line were retained and nucleotide changes in coding sequences were considered. Three genetically linked mutations in coding sequences on chromosome 1 were identified (Fig. 2A). A co-segregation analysis was conducted on individual seedlings of the segregating F2 population using cleaved amplified polymorphic sequence (CAPS) markers developed for the three identified SNPs. This analysis revealed that the mutation in *HISN2*, but not the other SNPs, is completely linked to the *sune82* phenotype.

**Fig. 2.**
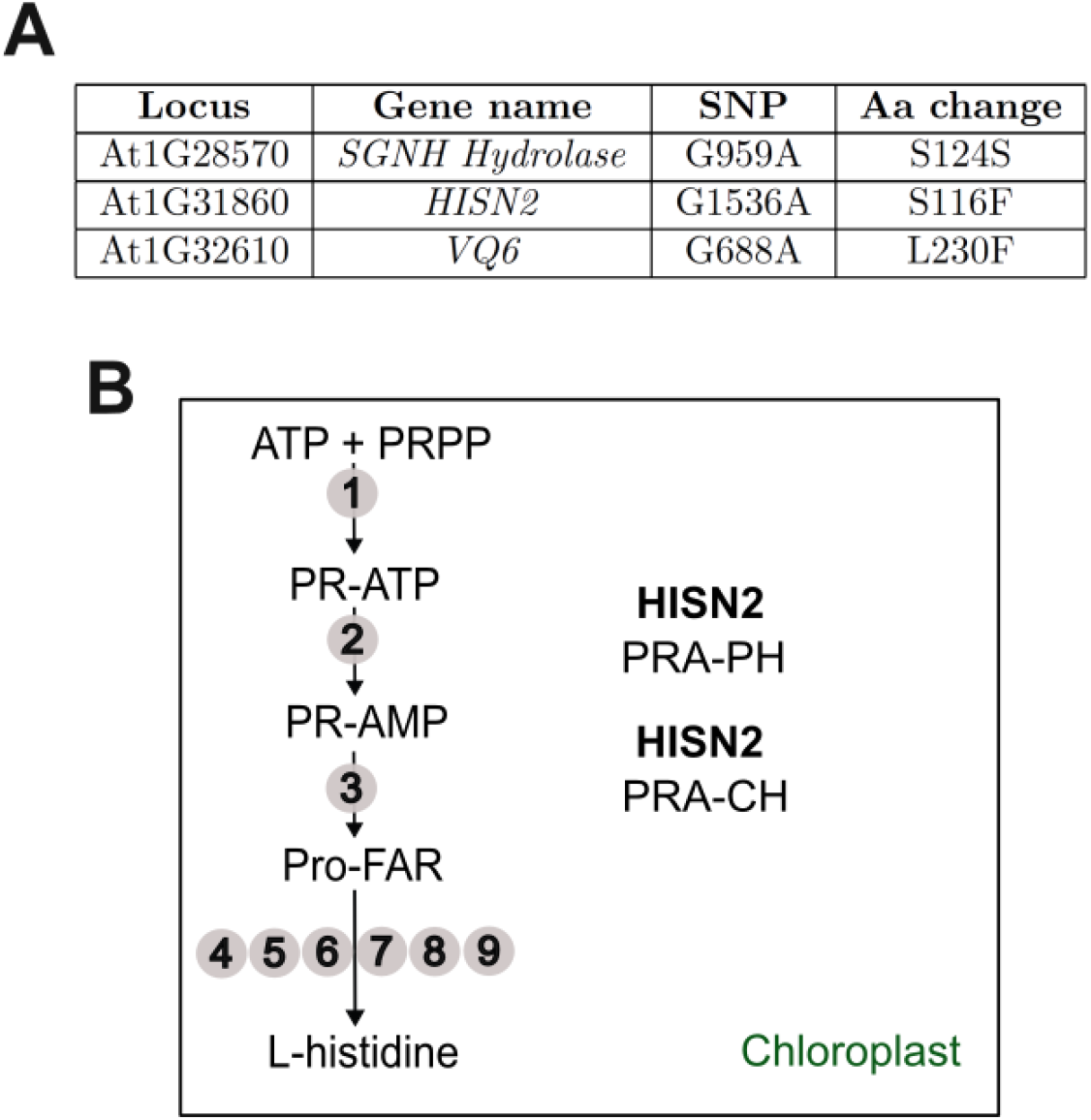
The *sune82* mutation affects the *HISN2* gene involved in His biosynthesis. (A) Three linked SNPs in coding sequences were identified in *LRX1ΔE14 sune82* compared to the non-mutagenized *LRX1ΔE14* line, of which the SNP in *HISN2* showed complete linkage. (B) HISN2 catalyzes the second and third steps of histidine biosynthesis in the chloroplast. Numbers indicate individual enzymatic steps in the pathway, those conducted by HISN2 are mentioned: No. 2, PRA-PH, phosphoribosyl-ATP pyrophosphatase, and No. 3, PRA-CH, phosphoribosyl-AMP cyclohydrolase.

*HISN2* (*AT1G31860*) encodes a bifunctional enzyme involved in His biosynthesis and is expressed in various tissues (Fig. 2B, Supplementary Fig. S2). HISN2 catalyzes the second and third steps in His biosynthesis in the chloroplast via two distinct domains: phosphoribosyl-ATP pyrophosphatase (PRA-PH) and phosphoribosyl-AMP cyclohydrolase (PRA-CH) (Fig. 2B). PRA-PH catalyzes the second step in which N1-5′-phosphoribosyl-ATP (PR-ATP) is hydrolyzed to N1-5′-phosphoribosyl-AMP (PR-AMP), while PRA-CH opens the adenine ring of PR-AMP to produce N1-[(5′-phosphoribosyl)formimino]-5-aminoimidazole-4-carboxamide- ribonucleotide (ProFAR). Six additional enzymes then convert this intermediate to histidine (Witek *et al*., 2021).

### Histidine biosynthesis is reduced in *sune82* plants

*LRX1ΔE14 sune82* plants contain a G-to-A substitution in *HISN2* causing a serine to phenylalanine residue substitution in the PRA-CH domain (Fig. 3A, Supplementary Fig. 3A). The crystal structure of HISN2 has recently been determined in *Medicago truncatula* (Witek *et al*., 2021). In the MtHISN2-AMP complex, PR-AMP interacts with the PRA-CH domain in a specific region formed by residues _107_WTKGETS_113_. Amino acid alignment of diverse species shows that this PR-AMP binding region is highly conserved throughout all domains of life (Fig. 3A). Remarkably, Witek *et al*. (2021) observed that substitution of the polar amino acid serine in this motif with alanine (MtS113A) reduces the activity of MtHISN2. Hence, the substitution of the corresponding position in the Arabidopsis HISN2 (AtS116F) in *sune82* could potentially reduce the activity of HISN2.

**Fig. 3.**
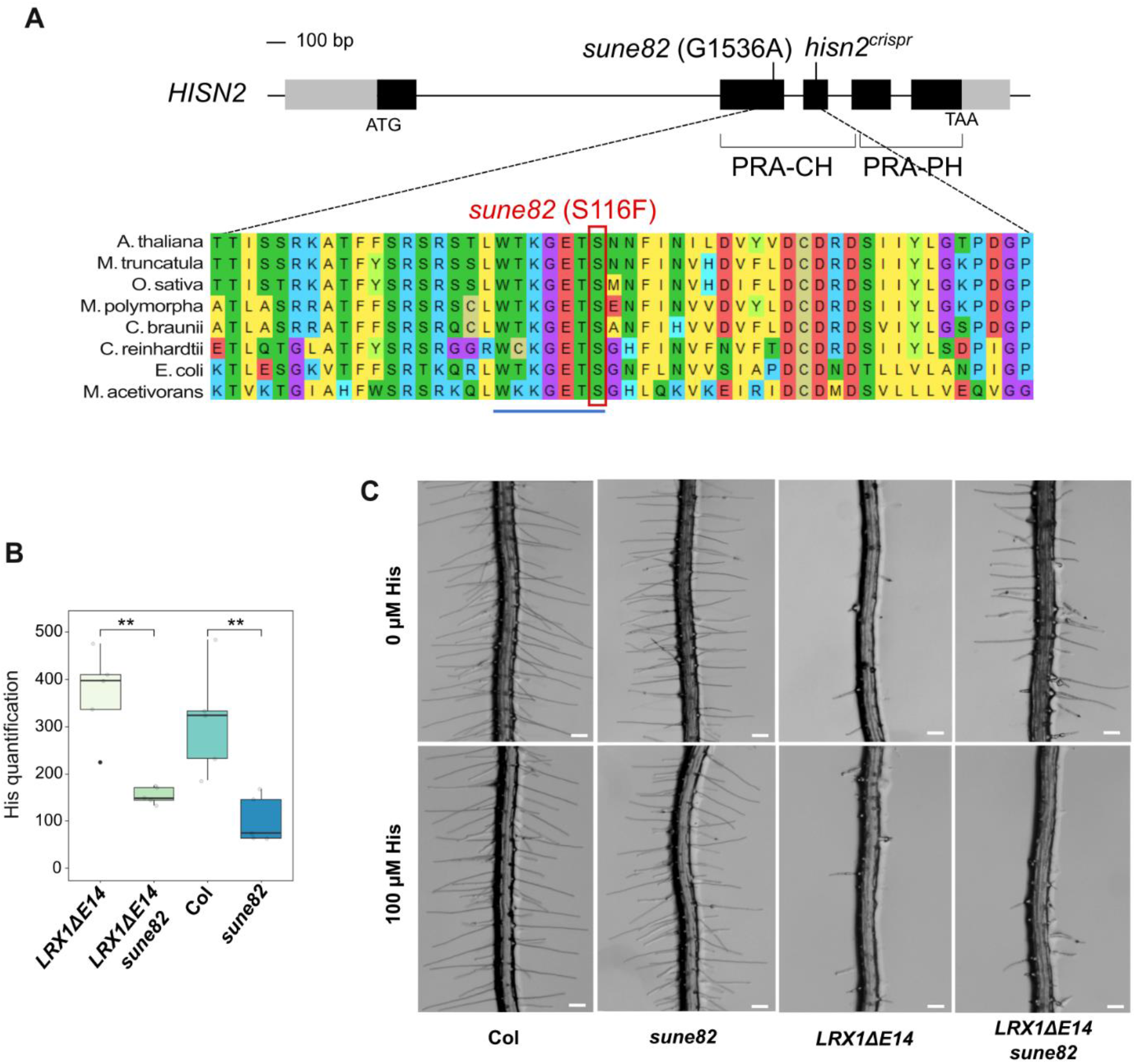
The *sune82* mutation changes the active site of *HISN2*, leading to a reduced histidine production. (A) Genomic structure of *HISN2* (AT1G31860) and amino acid alignment with its orthologs using MUSCLE in MEGA11. Grey boxes=UTR, black boxes=exons, lines=introns. PRA-CH: phosphoribosyl-AMP cyclohydrolase, PRA-PH: phosphoribosyl-ATP pyrophosphatase. *Arabidopsis thaliana, Medicago truncatula, Oryza sativa, Marchantia polymorpha, Chara braunii, Chlamydomonas reinhardtii, Escherichia coli, Methanosarcina acetivorans*. The Ser116 residue altered by the *sune82* mutation in boxed in red. (B) His levels were determined by LC-MS in 10-day-old seedlings. Arbitrary units show that His content is reduced by the *sune82* mutation. Asterisks on the graph indicate significant differences between genotypes (n=5, **P<0.01, unpaired *t*-test). Black line in the boxplots represents the median. (C) 5-day-old roots of Col, *LRX1ΔE14*, *LRX1ΔE14 sune82,* and *sune82* grown with or without 100 µM histidine. Bars= 200 μm.

To determine the impact of the *sune82* mutation on His production, His levels were determined by LC-MS in 10-day-old seedlings. *sune82* and *LRX1ΔE14 sune82* lines exhibited a 60% reduction in His content compared to wild-type Col and *LRX1ΔE14*, respectively (Fig. 3B). It is noteworthy that a comparable reduction was observed for the activity of HISN2_S113A_ vs HISN2_WT_ in *Medicago truncatula* (Witek *et al*., 2021), suggesting that the AtS116F substitution affects HISN2 activity to a similar degree.

We next wanted to confirm that the lack of His was responsible for the suppression of the *LRX1ΔE14* phenotype. *LRX1ΔE14* and *LRX1ΔE14 sune82* seedlings were germinated on media supplemented with or without 100 µM His. When grown on 100 µM His, the root hair phenotype of *LRX1ΔE14 sune82* seedlings was comparable to *LRX1ΔE14* (Fig. 3C), with shorter and burst root hairs, indicating that His supplementation can compensate for the *sune82* mutation. The short-root phenotype of *sune82* plants could also be suppressed by His supplementation (Supplementary Fig. S3B, Supplementary Fig. S3C), confirming that the reduced His content causes the dwarf phenotype and the suppression of *LRX1ΔE14*.

### A *HISN2* knock-out mutant is lethal

To further confirm that mutations in *HISN2* confer the suppressor phenotype in *LRX1ΔE14 sune82*, we generated a knock-out allele of *HISN2* in the *LRX1ΔE14* background using the CRISPR/Cas9 technology. A guide RNA targeting the third exon of *HISN2,* located in the PRA-CH coding domain, was used to obtain a *hisn2^crispr^* allele containing an A insertion, leading to a preliminary stop codon at the end of the PRA-CH domain (Supplementary Fig. S4A). It is known that His is essential for plant survival and that a complete blocking of His biosynthesis is embryo lethal (Muralla *et al*., 2007). When the progeny of a heterozygous *hisn2^crispr^* mutant was sown, 36% developed shorter roots and sequencing revealed these to be heterozygous for the *hisn2^crispr^* mutation (Supplementary Fig. S4). Importantly, heterozygous *hisn2^crispr^* seedlings showed suppression of the *LRX1ΔE14* root hair phenotype (Fig. 4), suggesting haplo-insufficiency (Deutschbauer *et al*., 2005). No homozygous *hisn2^crispr^* line could be identified among these seedlings. This data shows that *hisn2^crispr^* is a recessive lethal mutation and that *hisn2^crispr^* heterozygous plants likely have reduced HISN2 activity. To investigate this point further, seeds of the segregating population that did not germinate on regular MS were supplemented with 100 µM His. Under these conditions, germination was possible, and these seedlings revealed to be homozygous for *hisn2^crispr^* mutation. Additionally, the observed frequency of 5% homozygotes and 36% heterozygotes is below the expected value of 25% and 50%, respectively, if it were to follow Mendelian segregation. As seen with the segregation of *sune82* (see above), *HISN2* appears to influence the fertilization process. This confirms the importance of His synthesis for plant growth and development and demonstrates the lethality of a *hisn2* knock-out allele.

**Fig. 4.**
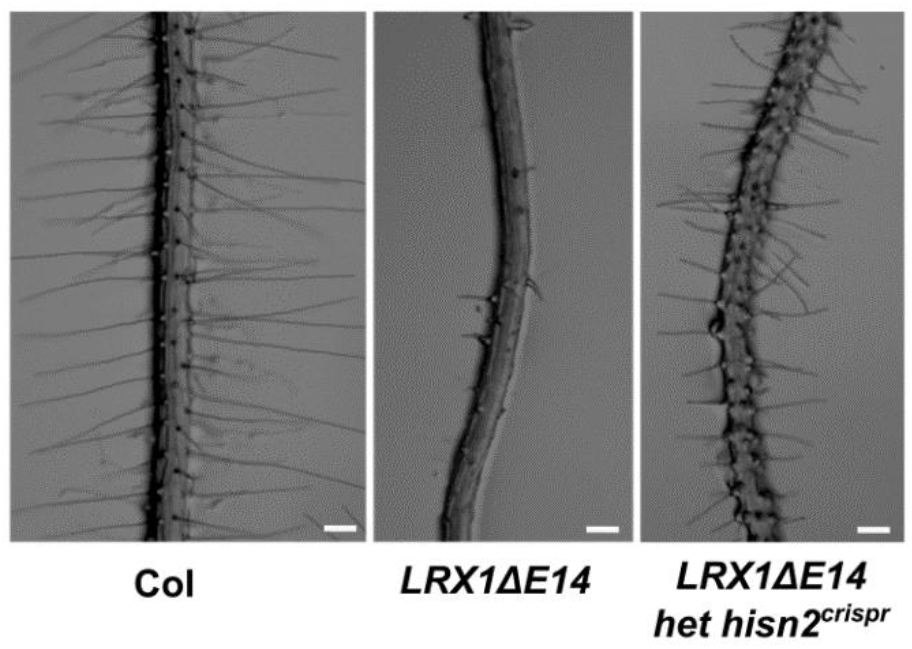
Heterozygous *hisn2^crispr^* seedlings suppress *LRX1ΔE14* root hair phenotype. 5-day-old roots of seedlings of wild-type Col, *LRX1ΔE14* and *LRX1ΔE14* heterozygous for the *hisn2^crispr^* allele reveal haplo-insufficiency in the latter and suppression of *LRX1ΔE14*. Bars= 200 μm.

### Reduced HISN2 activity leads to an altered TOR sensitivity

Histidine levels are reduced in the *sune82* mutant leading to a dwarf phenotype. Amino acid levels have been demonstrated to directly influence the target of rapamycin (TOR) network, which coordinates plant growth (Cao *et al*., 2019; Liu *et al*., 2021; Mallén-Ponce *et al*., 2022). In particular, His has been shown to activate TOR with moderate potency when exogenously supplied to inorganic nitrogen-starved seedlings (Song *et al*., 2022). Therefore, we hypothesized that the reduced production of histidine would affect TOR activity, leading to a change in plant growth. To first test whether TOR activity was impacted, we employed a specific TOR kinase inhibitor, AZD-8055, that targets the TOR kinase. Sensitivity to TOR inhibitors is frequently used to identify modulations in the TOR network that influence TOR kinase activity (Chan *et al*., 2000; Leiber *et al*., 2010; Barrada *et al*., 2019; Schaufelberger *et al*., 2019). Treatment of wild-type seedlings with AZD-8055 inhibits cell growth and root hair elongation (Montané and Menand, 2013). To determine whether *sune82* mutant has an altered sensitivity to AZD-8055 treatment, *sune82* and WT seedlings were germinated at different concentrations of AZD-8055. Already when applied at a low concentration (0.2 µM), AZD-8055 treatment decreased root hair length in Col and *sune82*, and the suppression of the *LRX1ΔE14* phenotype mediated by the *sune82* mutation was abolished (Fig. 5A). This indicates that TOR kinase activity is involved in the suppression of the *LRX1ΔE14* root hair phenotype.

**Fig. 5.**
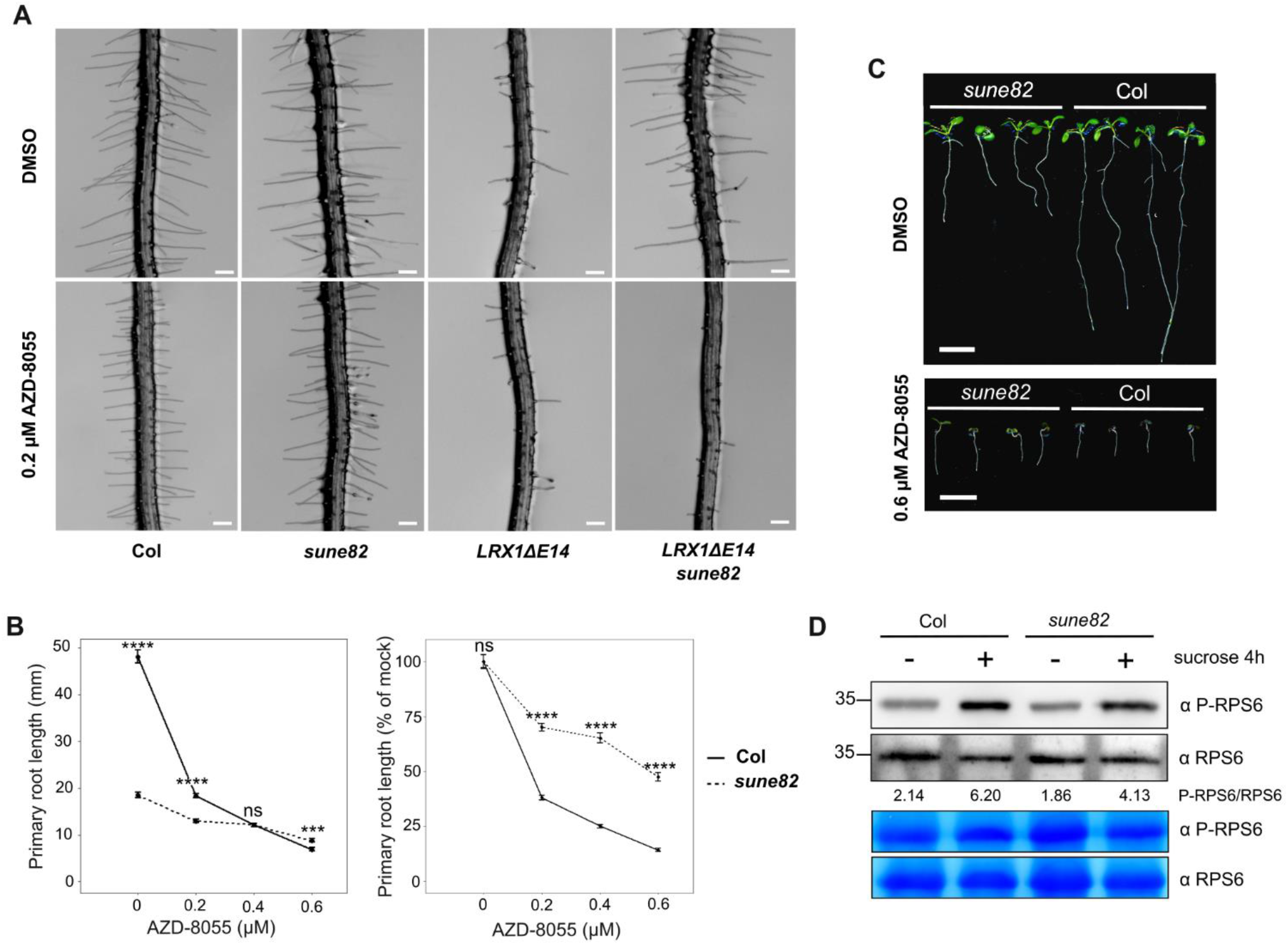
Reduced HISN2 activity leads to an altered TOR sensitivity. (A) 5-day-old roots of Col, *sune82, LRX1ΔE14*, and *LRX1ΔE14 sune82* grown with DMSO (mock) or 0.2 µM AZD-8055. Bars= 200 µm. (B) Primary root length quantification of 9-day-old seedlings grown at different concentrations of AZD-8055. First panel displays absolute primary root length (mm), second panel shows relative root length compared to mock conditions. Asterisks on the graph indicate significant differences between genotypes (n≥ 20, ****P<0.0001, *** P<0.001, ns P>0.05 and not significant, unpaired *t*-test). (C) 9-day-old seedlings grown on 0.6 µM AZD-8055. The primary root length of Col is significantly reduced, while that of *sune82* is almost unaffected. Bars= 1 cm. (D) For RPS6 analysis, six-day-old seedlings were transferred to sugar-free medium for 24 h and then either mock or sucrose (0.5%) treated for 4 h to induce RPS6 phosphorylation. 30 µg of whole-seedling extracts were subjected to immunoblot analysis with anti-phospho-RPS6 Ser240 or anti total RPS6 antibodies and ratios of anti-phospho-RPS6 Ser240 / anti-total RSP6 are indicated. Coomassie blue to ensure equal protein loading is also shown.

Next, the impact of AZD-8055 on primary root growth was assessed. *sune82* seedlings were strikingly less affected by AZD-8055 treatment than the wild type at all concentrations (Fig. 5B). At the concentration of 0.6 µM AZD-8055, *sune82* showed a decrease of around 50% in primary root length compared with mock conditions, whereas wild-type primary root length showed a decrease of 90% (Fig. 5B, C). *LRX1ΔE14 sune82* seedlings also presented a reduced AZD-8055 sensitivity compared to *LRX1ΔE14* regarding root length (Supplementary Fig. S5A, Supplementary Fig. S5B). Strikingly, growing seedlings for a longer period on 0.6 µM AZD-8055 revealed that *sune82* and *LRX1ΔE14 sune82* seedlings developed greener leaves and longer primary roots than the wild type and *LRX1ΔE14*, respectively (Supplementary Fig. S5C). These results indicate that TOR activity is affected in the *sune82* mutant. Interestingly, the TOR kinase activity seems to be activated differently among tissues: while *sune82* root hairs are sensitive to AZD-8055 similar to the wild type, most other tissues appear more resistant.

The TOR complex promotes cell proliferation and its inactivation leads to a reduction in meristem size and induces early differentiation (Ren *et al*., 2012; Cao *et al*., 2019). To determine whether *sune82* has an altered meristem size, root meristems of *sune82* and Col seedlings were examined using propidium iodide (PI) staining for visualization of the individual cells. *sune82* seedlings showed a reduction in meristem size and cell number, with a transition zone (TSZ), which separates dividing cells from differentiating cells into two functional domains, appearing closer to the root tip than in Col (Supplementary Fig. S5D, Supplementary Fig. S5E). This suggests that reduced cell proliferation in the meristematic region contributes to the reduced primary root length observed in seedlings containing the *sune82* mutation. Given the reduced sensitivity to the TOR inhibitor AZD-8055 and the short meristem size and primary root length of the *sune82* mutant, we investigated whether TOR activity was decreased at the seedling stage, using phosphorylation of ribosomal protein S6 (RPS6^S240^) as a readout. RPS6 is a downstream effector of TOR that is commonly used to measure TOR activation in Arabidopsis (Ren *et al*., 2012; Dobrenel *et al*., 2016*b*; Forzani *et al*., 2019; Mallén-Ponce *et al*., 2022). Antibodies against RPS6 and phosphorylated RPS6, respectively, that allow to assess phosphorylation levels, revealed no obvious difference between the wild type and *sune82* mutant (Fig. 5D). In addition, when TOR activity was induced by sucrose treatment, RPS6 phosphorylation increased to similar levels in *sune82* and wild-type seedlings. This result indicates that the RPS6 pathway is not downregulated in *sune82*. Taken together, these data indicate that the *sune82* phenotype is associated with an altered TOR network, but RPS6 phosphorylation seems not to be a major target of this altered activity.

### Alleles of *IPMS1* suppress *LRX1ΔE14*

The finding of a suppressor of *LRX1ΔE14* associated with the TOR pathway prompted us to investigate whether other known modifiers would also be found in the *sune* mutant collection. Mutations in *IPMS1*, an enzyme required for Leu biosynthesis, affect amino acid homeostasis and alter the TOR network (Cao *et al*., 2019; Schaufelberger *et al*., 2019). Consequently, the *IPMS1* locus of all *sune* mutants initially identified and subsequently confirmed was sequenced. One line, *sune106,* was found to have a C to T mutation in *IPMS1*, resulting in a Gly99Asp substitution in a highly conserved stretch of amino acids (Fig. 6, Supplementary Fig. S6A, Supplementary Fig. S6B). To confirm the causative effect of the mutation in *IPMS1*, an additional mutation was introduced in *IPMS1* in the *LRX1ΔE14* background using the CRISPR/Cas9 technology. A guide RNA targeting a sequence adjacent to the SNP of the *sune106* allele was utilized, and an *LRX1ΔE14* line homozygous for an insertion of an adenine leading to a premature stop codon was produced (Supplementary Fig. S6A, Supplementary Fig. S6C). This *ipms1^crispr^* line also showed suppression of the *LRX1ΔE14* root hair phenotype (Fig. 6), corroborating that the causative mutation in *sune106* is in *IPMS1*. Hence, modulating the TOR network by interfering with the Leu biosynthetic pathway also causes suppression of the *LRX1ΔE14* phenotype.

**Fig. 6.**
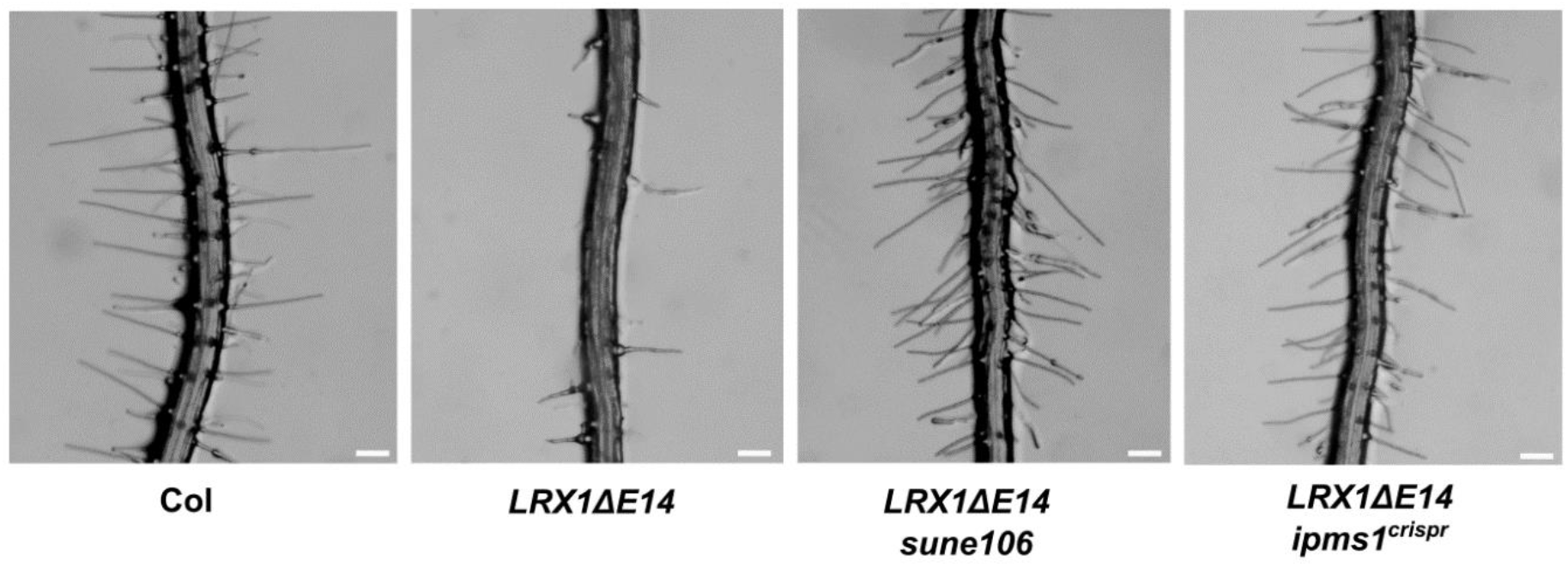
*sune106* suppresses *LRX1ΔE14* root hair phenotype. Five-day-old roots of Col, *LRX1ΔE14*, *LRX1ΔE14 sune106*, *LRX1ΔE14 ipms1^crispr^* show suppression of the *LRX1ΔE14* phenotype by mutations in *IPMS1*. Bars= 200 µm.

## DISCUSSION

### The *sune82* mutation leads to a partial activity of HISN2

His is an essential amino acid that is required for plant growth and reproduction (Mo *et al*., 2006; Muralla *et al*., 2007). In contrast to the majority of amino acids which are produced by enzymes encoded by multi-gene families, five of the eight His biosynthesis enzymes are encoded by single-copy genes in Arabidopsis (Muralla *et al*., 2007; Ingle, 2011). Knock-out mutations in most of these genes have therefore lethal effects (DeFraia and Leustek, 2004; Tzafrir *et al*., 2004; Muralla *et al*., 2007; Boavida *et al*., 2009; Petersen *et al*., 2010; Meinke, 2020). To date, two viable weak alleles of His biosynthesis genes have been identified in Arabidopsis. The *apg10* mutant carries a Val256Leu mutation in *HISN3* and exhibits a pale green phenotype in seedlings. Unlike embryo-lethal *hisn3* knock-out mutants, *apg10* plants gradually recover and are wild type-like in reproductive tissues (Noutoshi *et al*., 2005). The *hpa1* mutant presents an Ala69Thr substitution in *HISN6A*, which results in impaired root development in seedlings. Nevertheless, adult plants are indistinguishable from the wild type, potentially due to a gain in expression of its paralog *HISN6B* (Mo *et al*., 2006). Interestingly, *apg10* and *hpa1* have different effects on His content. The *apg10* mutation does not reduce free His content compared to WT, but does lead to a general increase in amino acid biosynthesis (Noutoshi *et al*., 2005). *hpa1* mutant seedlings exhibit a 30% reduction in free His content, but also show lower levels of free Asp, Lys, Arg, and Glu (Mo *et al*., 2006). Here, we characterized a novel His-deficient mutant, which presents a 60% decrease in free His content. The *sune82* mutant carries a weak allele of *HISN2* and exhibits altered development at both the seedling and adult stages. *sune82* displays an overall dwarf phenotype, with reduced primary root length and impaired fertilization. It is known that knock-out alleles of His biosynthesis genes cause embryonic lethality when homozygous and, in heterozygous plants, also affect gametophyte viability, resulting in reduced transmission of the mutant allele (Muralla *et al*., 2007). The *sune82* mutant exhibits a similar effect, with fertilization being significantly impaired, leading to a decrease in the frequency of homozygous mutants in the progeny of a heterozygous plant. This effect is weaker in *sune82* than in *hisn2^crispr^* mutant which phenocopies the *hisn2* knock-out mutants *hisn2-1* and *hisn2-2* (Muralla *et al*., 2007). Therefore, *sune82* represents the first viable *HISN* mutant identified that exhibits a significant reduction in His content and a mutant phenotype in most, if not all, tissues and developmental stages investigated.

The missense mutation Ser116Phe in *sune82* affects a residue localized in a highly conserved motif that is directly involved in AMP binding. This has been demonstrated in a detailed structural analysis of the HISN2 enzyme of *Medicago truncatula* (MtHISN2), where Ser113 corresponds to Ser116 of HISN2 of Arabidopsis (Fig. 3A). Interestingly, Ser113 of MtHISN2 is the sole residue in this motif that can be modified without completely losing enzymatic activity (Witek *et al*., 2021). The *sune82* mutant thus provides *in vivo* evidence that changing the polar Ser116 to a hydrophobic amino acid alters HISN2 activity, confirming the *in vitro* findings of Witek *et al*. (2021). Therefore, Ser116 in AtHISN2 may be one of the few residues of HISN2 that can result in an intermediate enzymatic activity and, consequently, to a significant but non-lethal growth phenotype.

### *sune8*2 can suppress the *LRX1ΔE14*-induced root hair growth defect in a TOR-dependent manner

The reduced His content induced by the *sune82* mutation not only leads to a dwarf phenotype but also results in an enhanced root hair development and suppression of the dominant-negative effect induced by *LRX1ΔE14*. Similarly, the alteration in amino acid homeostasis mediated by the *sune106* mutation by affecting IPMS1, suppresses the *LRX1ΔE14* root hair phenotype. As in yeast and in animal cells, the plant TOR network senses nutrient availability to adjust cell growth (Robaglia *et al*., 2012; Liu *et al*., 2021; Li *et al*., 2023). The use of *TOR* RNAi lines and treatments with inhibitors of the TOR kinase has demonstrated that TOR influences the accumulation of sugars such as raffinose, amino acids, and a number of secondary metabolites (Moreau *et al*., 2012; Ren *et al*., 2012) and in this way regulates cell wall remodeling and cell growth (Calderan-Rodrigues and Caldana, 2024). Treatment with the TOR inhibitor AZD-8055 to WT seedlings reduces root hair development. A comparable effect was observed in *LRX1ΔE14 sune82* seedlings grown in the presence of AZD-8055, which developed an *LRX1ΔE14* phenotype. This indicates that TOR is involved in the suppression of *LRX1ΔE14* mediated by the *sune82* mutation. Interestingly, in other tissues, seedlings were more resistant to this inhibitor than the wild type: root growth inhibition was less pronounced and cotyledon leaves were greener. Thus, while TOR function is demonstrably impaired in primary root development, it remains operational during root hair formation. It is conceivable that lack of histidine causes TOR to be up-regulated in root hairs since these are essential for effective uptake of nutrients from the soil (Miguel *et al*., 2015; Morris *et al*., 2017) and inhibition of translational activity was shown to activate TOR (Watanabe-Asano *et al*., 2014). Moreover, it is known that in low nitrate conditions root hair growth is stimulated and this process is TOR-dependent (Pacheco *et al*., 2023).

By contrast, TOR is likely down-regulated in other tissues due to the lack of His (Heinemann and Hildebrandt, 2021). Indeed, the comparable phenotype of mutations in *IPMS1* involved in Leu biosynthesis supports this view (Schaufelberger *et al*., 2019 and data shown above). Propidium staining revealed that *sune82* roots are affected early on in the differentiation of meristematic cells, resulting in a transition zone appearing closer to the root tip than in wild-type seedlings. This is consistent with the previous observation that TOR influences plant growth by inducing early differentiation of meristematic cells (Montané and Menand, 2013). This similarity of root development in *sune82* and the AZD-8055-treated wild type possibly explains the reduced sensitivity of *sune82* to AZD-8055.

Our analysis on the phosphorylation dynamics of RPS6, a ribosomal protein phosphorylated by the S6 kinase 1 (S6K1) that is a direct target of the TOR kinase (Mahfouz *et al*., 2006), did not reveal significant changes in phosphorylation in *sune82* and the wild type. Upon addition of sucrose, RPS6 is readily phosphorylated as previously shown (Dobrenel *et al*., 2016*b*), but again, no obvious difference was observed between the wild type and *sune82*. It is plausible that the alterations in the *sune82* line are rather subtle, while changes in RPS6 phosphorylation are mainly seen when the TOR network is strongly influenced, as exemplified by the treatment with the TOR kinase inhibitor AZD-8055 or by silencing TOR expression (Dobrenel *et al*., 2016*b*). The TOR kinase is involved in numerous processes (Ingargiola *et al*., 2020), and it is also possible that numerous downstream effectors are subtly altered, the sum of which results in the observed growth alteration in *sune82*. Alternatively, RPS6 phosphorylation might vary in different cell types but single-cell type analyses are technically difficult to perform.

The CrRLK1L receptor kinase FER, was recently shown to directly phosphorylate and activate the TOR kinase in a RALF (Rapid ALkalinization Factors)-enhanced manner (Pearce *et al*., 2001; Song *et al*., 2022; Pacheco *et al*., 2023; Liu *et al*., 2024). This finding establishes a link between the TOR network and a major cell growth control machinery. LRX are high-affinity receptors of RALF peptides (Mecchia *et al*., 2017; Dünser *et al*., 2019; Moussu *et al*., 2020) and function in conjunction with FER to regulate a number of processes, including development (Dünser *et al*., 2019; Herger *et al*., 2019; Gronnier *et al*., 2022). This provides an explanation why LRX1-related phenotypes are influenced by a modification of the TOR network (Baumberger, 2001; Leiber *et al*., 2010; Schaufelberger *et al*., 2019). In this study, the *sune82* mutant was found to suppress the dominant-negative phenotype of *LRX1ΔE14*. The LRX1ΔE14 protein lacks the extensin domain that anchors LRX proteins in the cell wall (Rubinstein *et al*., 1995; Baumberger, 2001), which results in its inability to become insolubilized in the cell wall (Ringli, 2010). LRXs were recently shown to influence the compaction of pectin, a major component of the cell wall (Moussu *et al*., 2023; Schoenaers *et al*., 2024). Hence, LRX1ΔE14 might form inadequate LRX-RALF-pectin connections, which interferes with proper structuring of the cell wall, resulting in defective root hairs. These mal-forming cell wall structures might be perceived by the FER transmembrane receptor kinase that transmits corresponding information to the TOR network (Song *et al*., 2022; Pacheco *et al*., 2023), establishing a link between LRX1/ LRX1ΔE14 activity and the TOR network.

In conclusion, we have identified a weak allele in a gene involved in His biosynthesis, that results in a mutant with consistent alterations in plant development. In contrast to other mutants affected in this pathway, the *sune82* mutant phenotype is neither lethal nor restricted to a short period during the plant life cycle. This weak *HISN2* allele *sune82* allowed to establish a link between an *LRX1ΔE14*-induced root hair defect and the TOR network, which is likely affected due to the reduced availability of His. Cell wall integrity sensing mechanisms directly linking to the TOR kinase may be a plausible explanation for the observed suppression of the *LRX1ΔE14*-induced root hair defect by *sune82*, but the molecular details on a higher-resolution on the subcellular level remain to be investigated.

## SUPPLEMENTARY DATA

The following supplementary data are available at JXB online.

Table S1. Primers used in this study.

Fig. S1. *sune82* plants display a dwarf phenotype.

Fig. S2. Expression pattern of *HISN2*.

Fig. S3. Histidine supplementation can complement *LRX1ΔE14 sune82* root phenotype.

Fig. S4. Homozygous *hisn2^crispr^* mutants are lethal.

Fig. S5. The *sune82* mutation reduces sensitivity to the TOR inhibitor AZD-8055 and results in a shorter meristem.

Fig. S6. *sune106* and *ipms1* crispr mutations are in the first exon of *IPMS1*.

## ACKNOWLEDGEMENTS

This work was supported by the Swiss National Science Foundation, grants No 31003A_166577/1 and 310030_192495 to CR. Furthermore, we thank the infrastructure team at DPMB for continuous support with growth facilities.

## AUTHOR CONTRIBUTION

Conceptualization: CR; methodology: AH, AG, AGS, CM; formal analysis: AG, CL, AH, DR, AGS, XH, MS; resources: CR, TW; data curation: AG, AGS, TW; AG, CL, CR; writing - original draft: AG, CR; writing - review & editing: all authors; visualization; AG, AH, CL, CR, LB; supervision: CR, TW, LB; funding acquisition: CR.

## DATA AVAILABILITY

All data are made available upon publication in a publicly accessible repository.

## Abbreviations

EMS, ethyl methanesulfonate; FER, FERONIA; HISN2, histidine biosynthesis 2; IPMS1, isopropylmalate synthase 1; LRX, leucine-rich repeat extensin; MS, Murashige and Skoog; PRA-CH, phosphoribosyl-AMP cyclohydrolase; PRA-PH, phosphoribosyl-ATP pyrophosphatase; PI, propidium iodide; RALF, rapid alkalinization factor; RPS6, ribosomal protein S6; *sune*, *suppressor of dominant-negative LRX1ΔE14;* TOR, target of rapamycin.

